# An open-source platform to distribute and interpret data from multiplexed assays of variant effect

**DOI:** 10.1101/555797

**Authors:** Daniel Esposito, Jochen Weile, Jay Shendure, Lea M Starita, Anthony T Papenfuss, Frederick P Roth, Douglas M Fowler, Alan F Rubin

## Abstract

Multiplex Assays of Variant Effect (MAVEs), such as deep mutational scans and massively parallel reporter assays, test thousands of sequence variants in a single experiment. Despite the importance of MAVE data for basic and clinical research, there is no standard resource for their discovery and distribution. Here we present MaveDB, a public repository for large-scale measurements of sequence variant impact, designed for interoperability with applications to interpret these datasets. We also describe the first of these applications, MaveVis, which retrieves, visualizes, and contextualizes variant effect maps. Together, the database and applications will empower the community to mine these powerful datasets.

## Background

Experimentally interrogating the effects of genetic variation has helped reveal the mechanisms by which genes function and facilitate an understanding of the clinical consequences of human genetic variation. Multiplex Assays of Variant Effect (MAVEs) leverage high-throughput DNA sequencing to greatly increase the scale at which variants can be experimentally investigated [1–3]. A MAVE yields a set of scores that describe the functional effect of thousands to tens of thousands of variants of a coding sequence, promoter, enhancer or other genetic element relative to a reference sequence. MAVEs are being adopted rapidly for both basic research and clinical applications [4]. As a consequence, the total number of variants with functional data generated by MAVEs was predicted to surpass 200,000 by the end of 2018 [3], which exceeds the number of classified missense variants available in ClinVar [5].

These large-scale variant effect maps are yielding insights into protein function, structure, and evolution [6–10]; exploring gene regulation and promoter function [11–13]; improving computational variant effect prediction [14, 15]; and guiding variant interpretation in the clinic [16–20]. However, the impact of variant effect maps has been limited by shortcomings in data availability, dissemination, and discoverability. Many publications describing large-scale variant effect mapping do not provide variant effect scores for all variants that were assayed. When variant effect scores are provided, they are often accessible only as a supplementary table or via a bespoke web interface [18, 19, 21] leading to a proliferation of inconsistent formats. Some publications, instead of including variant effect scores, deposit the associated high-throughput DNA sequencing data in the Sequence Read Archive or Gene Expression Omnibus [22, 23]. This raw data can be used to reconstruct variant effect scores, but accurately replicating the original analysis can be non-trivial. While databases of variant effect information exist, they are typically designed for a specific application [24–26] or a specific group of target genes [27–29]. Larger and more general databases can sometimes contain variant effect data [30, 31], but these platforms were not developed with large-scale variant effect maps in mind, so valuable context for the variant effect scores and associated metadata may be lost. Furthermore, most existing resources lack support for noncoding targets entirely.

To overcome these challenges and facilitate future advances, we are establishing an open-source platform for MAVE resources. The foundation is MaveDB, a central repository that allows researchers to store and publish processed MAVE datasets, associated metadata, and linked raw data using a machine readable, standardized, and searchable format. An easy-to-use web interface maximizes the impact and usefulness of researchers’ work by making data readily accessible to the whole community, whether for clinical applications, meta-analysis, or reanalysis as computational techniques are refined.

This platform is designed to allow additional applications to communicate directly with MaveDB. The first of potentially many such applications, MaveVis, visualizes and provides context to protein variant effect maps by generating heatmaps and automatically integrating them with secondary structure, surface accessibility, interaction interfaces, and conservation data.

## Organization and content of MaveDB

To capture the structure of real-world study designs, MaveDB is organized hierarchically into score sets, experiments, and experiment sets (**Figure 1**). Score sets, the most basic unit of organization, contain the variant effect scores and additional metadata such as target sequence information and detailed methods. Each variant effect score is a numeric value. Optional data columns containing values related to each variant effect score such as variant counts and measures of uncertainty can also be included and named by the user.

**Figure 1:**
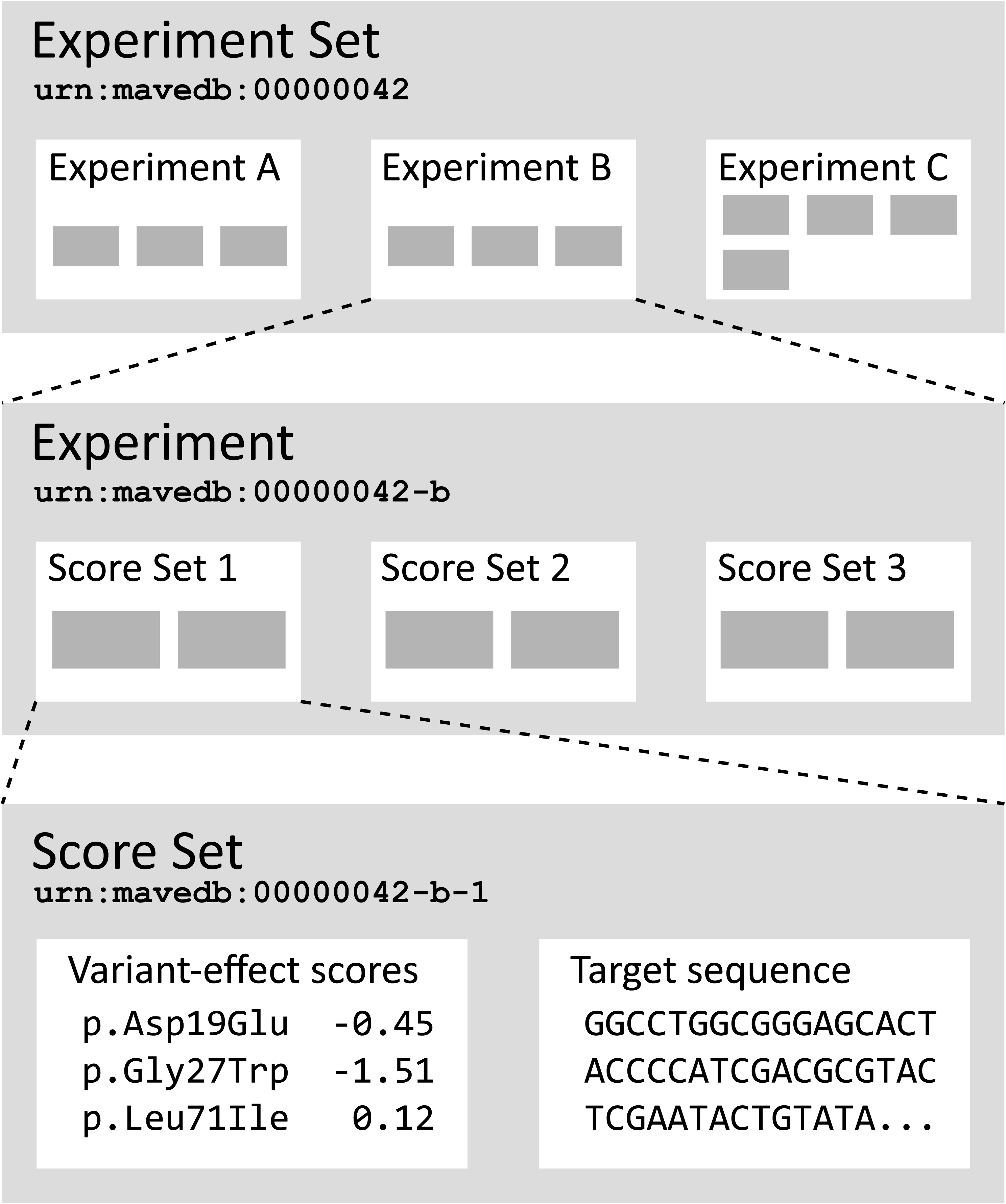
Relationships between MaveDB entries. MaveDB has three main entry types arranged in a hierarchical structure. The URN accession numbers shown for each example entity reflect these relationships. Additional metadata fields for score set and experiment entries are listed in **Table 1**.

Most experimental designs in MaveDB involve multiple score sets. For example, protein MAVEs commonly have one score set for nucleotide variants and another for amino acid variants [32]. Experiments with tiled designs [33, 34] can have score sets for each tile, and experiments with multiple distinct reference sequences [35] can have score sets for each reference sequence [35][40]. In addition, we envision that reanalysis and renormalization of existing datasets using updated methods will be commonplace [14, 15, 36, 37]. By grouping all analyses of a single raw dataset under one experiment, MaveDB ensures that the number of assays performed on each target sequence can be tracked accurately.

**Table 1:** MaveDB metadata fields

| Field name | Experiment | Score Set | Type | Searchable | Link |
| --- | --- | --- | --- | --- | --- |
| Keyword | ✓ | ✓ | String | ✓ |  |
| Abstract | ✓ | ✓ | Markdown | ✓ |  |
| Method | ✓ | ✓ | Markdown | ✓ |  |
| Short description | ✓ | ✓ | String | ✓ |  |
| Title | ✓ | ✓ | String | ✓ |  |
| PubMed ID | ✓ | ✓ | Accession | ✓ | ✓ |
| DOI | ✓ | ✓ | Accession | ✓ | ✓ |
| SRA accession | ✓ |  | Accession | ✓ | ✓ |
| RefSeq accession |  | ✓ | Accession | ✓ | ✓ |
| Ensembl accession |  | ✓ | Accession | ✓ | ✓ |
| UniProt accession |  | ✓ | Accession | ✓ | ✓ |
| Created by | ✓ | ✓ | Contributor | ✓ | ✓ |
| Last modified by | ✓ | ✓ | Contributor | ✓ | ✓ |
| Creation date | ✓ | ✓ | DateTime | ✓ |  |
| Modification date | ✓ | ✓ | DateTime | ✓ |  |
| Publication date | ✓ | ✓ | DateTime | ✓ |  |
| Licence |  | ✓ | Licence | ✓ | ✓ |
| Has replacement |  | ✓ | Boolean | ✓ |  |

Each experiment describes one or more analyses arising from a single MAVE, including any technical and biological replicates. In addition to links to score sets, experiments contain metadata including methodological details, links to raw data, and associated publications (**Table 1**), but no variant score information. Experiment sets contain one or more related experiments, for example multiple MAVEs performed on the same target sequence under different conditions or multiple experiments from the same publication.

MaveDB currently contains over one million variant effect scores across 39 unique targets. We welcome both new and previously described datasets from the community and have implemented a conversion tool, mavedb-convert, for datasets generated by Enrich [38], Enrich2 [37], and EMPIRIC [39] (see Availability section).

## Utility of MaveDB

### Accessing datasets

MaveDB can be accessed through a standard web browser that allows users to explore by keyword, target gene, or organism. Alternatively, the advanced search function allows users to query all metadata fields, including the full text of methods and abstracts. Complete sets of variant effect scores and related values can be downloaded from any score set page in comma-separated value format. These files can be parsed easily in most scientific programming environments or imported into spreadsheet applications.

The advanced search function is also accessible programmatically through the REST API (Representational State Transfer Application Programming Interface). The API returns structured data, including full score sets and metadata, in JSON format, suitable for deserialization by most programming languages. Users of the R programming environment [40] can access MaveDB’s REST API using the rapimave library, which also includes a suite of exploration, searching, parsing and filtering functions (see Availability section).

### Creating new entries

Typically, a user starts by creating an experiment. The experiment can be added to an existing experiment set if desired, otherwise a new one will be created automatically. The user provides a description of the assay used to generate the raw data, adds links to the raw data if available, and can then add contributors. After the experiment is created, the user creates one or more associated score sets. Here, the user enters required information about the target such as its name and sequence, and also describes the methods used to calculate the variant effect scores from the raw data. Variant effect scores and optional counts files are then uploaded via the web interface and validated by the server.

### Publishing datasets

When first created, score sets, experiments, and experiment sets are private and have temporary accession numbers. Private entries are only viewable by their contributors and all values may be modified. Private entries can be accessed through the API by providing a contributor’s private access token generated on the contributor’s user profile page.

Completed private score sets can be published, making the score set publicly viewable. Publication creates a stable accession number and freezes the target sequence and variant effect score data, ensuring that all subsequent analyses based on the data are recomputable. Associated experiment and experiment sets are also published automatically if they are still private. Users may continue to edit some metadata such as the methods and description after publication.

Published scores cannot be changed, but in case a correction is necessary, MaveDB allows score sets to be deprecated when creating a replacement. Users browsing MaveDB will only see the most recent version, but deprecated score sets will remain available by accession number to ensure that previous analyses are reproducible.

### Contributor permissions

MaveDB supports three contributor roles: administrator, editor and viewer. Administrators can add or remove contributors, modify entries, and publish score sets. Editors can modify entries but cannot affect the contributor list or make entries public. Viewers can see their private entries in the database but cannot change them.

All three roles appear in the contributor list with no visual distinction between them, and administrators can continue to change the contributor list for each score set or experiment after publication. Since score sets and experiments have independent contributor lists, MaveDB maintains clear attribution when datasets are reanalyzed.

### Data licensing

Administrators may select one of several Creative Commons licenses for each score set [41–43] and additional licensing options may be added in response to user requests. The license information is included as score set metadata and as part of the header of each downloaded file. Administrators can relicense after publication, although users who download under a more permissive license would not be subject to a more restrictive license.

## Visualizing variant effect maps with MaveVis

The MaveVis application allows users to quickly visualize score sets retrieved directly from MaveDB. One example of MaveVis output for a variant effect map of the protein SUMO1 [44] is shown in **Figure 2**. Score sets are rendered as heatmaps with additional tracks representing integrated structural and conservation information from PDB [45] and UniprotKB [31]. The heatmap shows all possible amino acid changes at each protein sequence position, with colors reflecting the variant effect scores. The color scale is automatically calibrated based on the scores of reference and null alleles in the dataset or set manually by the user. Error bars are drawn directly on the heatmap fields to represent the measurement error provided in the score set, if present. Additional tracks show burial in protein interaction interfaces, residue-specific solvent accessibility, protein secondary structure, and sequence conservation.

**Figure 2:**
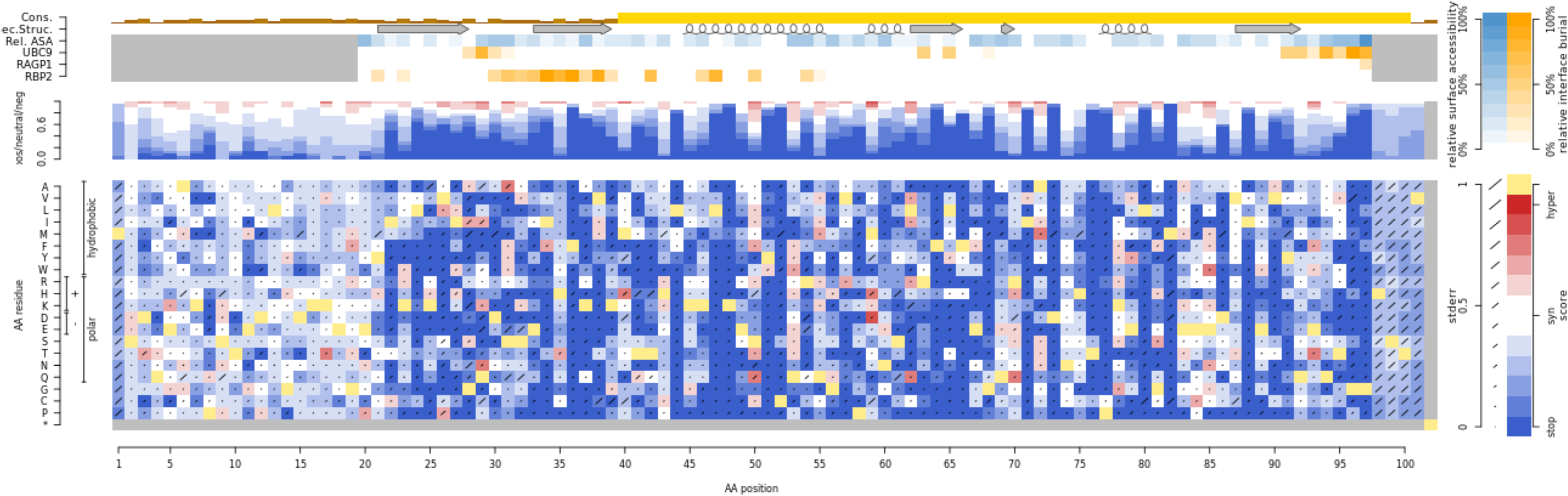
Heatmap for the SUMO1 MAVE dataset rendered by MaveVis. The x-axis iterates over amino acid positions in the protein, while the y-axis lists all possible amino acid changes organized by their physicochemical properties. The heatmap color reflects the variant effect score, with blue being as deleterious as a full deletion, white being equivalent to the reference allele, and red representing a stronger phenotype than the reference residue at that position. Yellow cells indicate the reference amino acid at each position. Error bars represent standard error of the mean. The stacked bars above the heatmap represent the relative frequencies for each phenotype bin of corresponding color at each position. Additional tracks show data integrated from other databases: orange heatmap tracks represent burial in protein interaction interfaces, the steel blue heatmap track represents solvent accessibility, the arrows and spirals correspond to secondary structure, and the yellow bar chart at the top shows sequence conservation.

### Accessing MaveVis

MaveVis is hosted at http://varianteffect.org, a portal for applications built on MaveDB. Users can follow the MaveVis link on each MaveDB score set page or navigate directly to http://vis.varianteffect.org and search for datasets. Once a score set is selected, the corresponding UniProt accession from MaveDB is suggested when available. MaveVis automatically presents potentially relevant PDB structures for the selected protein that overlap with the score set target sequence, allowing users to select which structures to include in the visualization. The resulting plot can be downloaded in PNG, PDF or SVG format.

In addition to the web interface, MaveVis also exists as an R package for local use (see Availability section). The R package provides direct access to both the visualization and underlying data integration functions, making it easy to automatically compile structural and conservation feature tables for individual proteins.

### Interaction with MaveDB

The MaveVis server automatically synchronizes with MaveDB at regular intervals via its API, caching any new score sets, automatically obtaining relevant PDB and UniProt data, and pre-calculating partial results for a more responsive user experience. MaveVis also exposes its own API, allowing it to be used within more complex workflows.

To facilitate communication between MaveVis and MaveDB, we developed an R package, hgvsParseR, to parse or assemble HGVS [46] strings that describe alleles (see Availability section). In addition to its utility for visualizing variant effect maps, we expect that this package will be generally useful for working with data from ClinVar [5], gnomAD [47], and other important sequence variation resources.

## Conclusions

MaveDB is the foundation of an open-source platform for the collection, distribution, and analysis of variant effect maps. Designed to be flexible and extensible, the MaveDB repository can accommodate data from diverse target sequences and experimental methods as the field evolves. Using MaveDB to combine variant effect data with external contextual information, MaveVis is the first application built on this resource. We envision developing additional applications such as tertiary structure analysis, automatic imputation of missing values in variant effect maps [48], and a broadly-applicable dashboard to assist dataset interpretation.

MaveDB, MaveVis, and Enrich2 simplify, standardize, and democratize MAVE data analysis. These tools are the beginnings of a community-driven, open-source platform that allows researchers to explore these comprehensive datasets. The impact of each dataset will continue to increase as the number of assayed variants grows, contributing to a more complete understanding of genetic variation and sequence function.

## Methods and implementation

MaveDB is implemented in Python using the Django Python Web framework [49, 50]. The relational database backend is PostgreSQL [51]. Asynchronous tasks such as handling file uploads and sending emails are managed using RabbitMQ and Celery [52, 53]. Variant score and count data are stored using PostgreSQL JSONField objects, which offer additional flexibility for storing arbitrarily-named data columns compared to a more traditional relational database design. Database accession numbers for publicly accessible entries are assigned in URN (Universal Resource Name) format [54] and their structure reflects MaveDB’s hierarchy (**Figure 1**).

Differences between each variant sequence and the target sequence are described using HGVS format [46]. MaveDB supports variant strings that describe substitutions or small indels in most sequence contexts.

Contributors are authenticated using their ORCID iD via the OAuth2 protocol [55, 56]. Consequently, an individual must have an ORCID iD to be named as a contributor to a MaveDB dataset. Users do not need to log in to browse or download publicly available data. MaveDB allows users to provide a private contact email address if they want to be contacted by administrators or receive alerts, but all other details are pulled from their public ORCID record.

Abstract and methods sections support Markdown [57] blocks for formatted text with support for mathematical notation. Markdown blocks are rendered to HTML using Pandoc [58].

MaveVis is implemented using R [40] and Docker [59]. Surface accessibility and interface burial are calculated using FreeSasa [60]. Secondary Structure is calculated using DSSP [61]. Conservation tracks are calculated using the AMAS algorithm [62], based on multiple alignments computed using ClustalOmega [63] for the appropriate UniRef90 set of orthologous proteins with at least 90% sequence identity from UniProtKB [31].

## Declarations

### Availability of data and material

MaveDB is hosted at https://www.mavedb.org/

MaveVis is hosted at http://vis.varianteffect.org/

Source code for all websites, tools, and packages is available on GitHub at https://github.com/VariantEffect

- MaveDB: https://github.com/VariantEffect/mavedb
- MaveVis: https://github.com/VariantEffect/mavevis
- mavedb-convert: https://github.com/VariantEffect/mavedb-convert
- rapimave: https://github.com/VariantEffect/rapimave
- hgvsParseR: https://github.com/VariantEffect/hgvsParseR

A pre-compiled docker image for MaveVis is also available on DockerHub at https://hub.docker.com/r/jweile/mavevis/

### Ethics approval and consent to participate

Not applicable.

### Authors’ contributions

A.F.R., D.M.F., and F.P.R. conceived of the project. D.E. and A.F.R. built the database and its web interface. J.W. built the MaveVis application. D.E. and J.W. built the APIs. All authors wrote the manuscript and approved the final version.

## Acknowledgements

We would like to acknowledge Bernie Pope, Peter Georgeson, Nick Moore, Matthew Wakefield, and Dan Bolon for helpful advice and guidance.

---

**Supplemental Figure 1:** UML (Unified Markup Language) diagram of the complete MaveDB schema in PDF format. The diagram was generated using the Django Extensions package and visualized using Graphviz [64, 65].

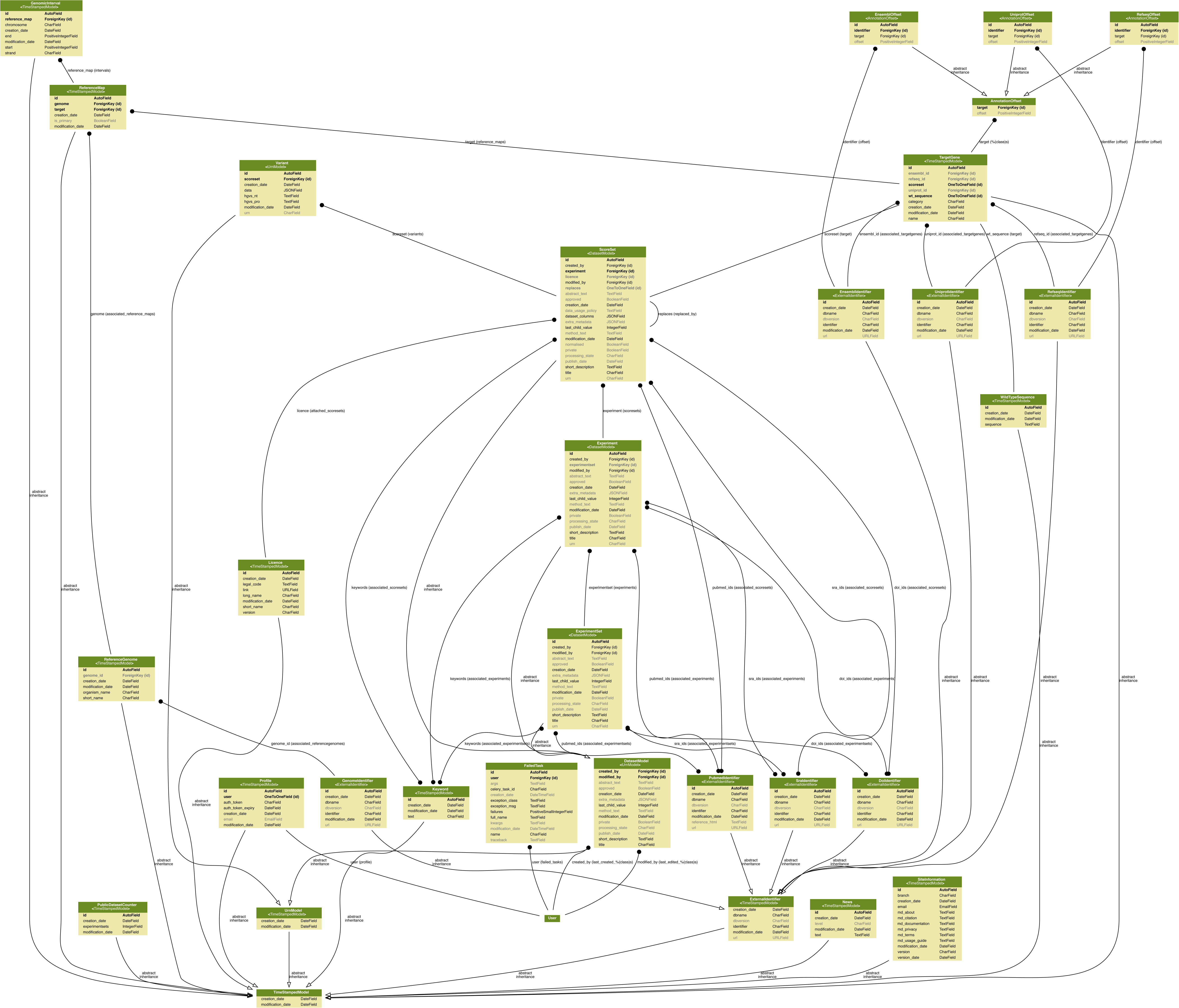

